# Dehydration Adaptation of the Adult Rat Modifies Brain Microtubule Electrical Activity of the Osmosensitive Supraoptic Nucleus

**DOI:** 10.1101/2025.11.18.688944

**Authors:** Horacio F. Cantiello, Cintia Y. Porcari, Virginia H. Albarracín, David Murphy, Andre S. Mecawi, María del Rocío Cantero, Andrea Godino

## Abstract

Brain microtubules (MTs) are important cytoskeletal structures in neurons that generate electrical oscillations in the frequency range of mammalian brain waves. However, the functional role of MT oscillations in the physiology of brain function remains unknown. Acute dehydration (DH) induced by water restriction results in changes in osmolality that require the modification of osmosensitive neuronal activity in the regions of the CNS involved in osmoregulation, in particular magnocellular neurons from the hypothalamic supraoptic nucleus (SON), responsible for the release of vasopressin and implicate osmosensor channels, associated with an intricate mesh of MTs. In the present study, we evaluated the effect of 24 h water deprivation (DH, dehydrated group) on the MT-based cytoskeleton of adult Wistar rats, as compared to a control group with free access to water (Control, euhydrated group). We obtained local field potentials (LFP) from isolated SON and cortex (CTX) brain tissue, observing spontaneous oscillations under both conditions. The electrical oscillations of the brain tissue from DH animals had different amplitudes and frequency peaks as compared to controls. A frequency domain spectral analysis of the time records indicated specific, SON-associated MT energy contributions of the frequency peaks in the challenged group, most particularly a dramatic increase in the 10-20 Hz range, and a statistically significant reduction in the high frequency 83-100 Hz region, not observed in the CTX samples. The data indicate that brain MTs respond to dehydration to produce electrical oscillations with distinct properties in the areas of the mammalian brain that specifically contribute to the homeostatic response. The present study provides the first evidence for a novel physiological mechanism associated with the electrical activity of the neuronal cytoskeleton and points to its possible involvement in the brain osmoregulatory mechanism.

## Introduction

The balance of body water and sodium levels is intricately regulated to prevent abnormal cardiovascular function and the onset of hypotension or hypertension^1,2^. Acute dehydration (DH), such as in a water deprivation condition, drives an elevation in osmolarity and a reduction in both intracellular and extracellular fluid compartments. In response, the neuropeptide hormones vasopressin (AVP) and oxytocin (OXT) are swiftly released from the posterior pituitary axon terminals of hypothalamic magnocellular neurons (MCNs) mainly located in the supraoptic nucleus (SON) and paraventricular nucleus (PVN) to restore plasma osmolarity through diuresis inhibition and natriuresis stimulation. The MCNs are inherently sensitive to osmotic changes and serve as secondary brain osmosensors, crucial for regulating thirst and AVP responses during dehydration^3-6^. Osmosensory transduction in osmosensitive neurons (ONs) is dependent, et least in part, by the N-terminal variant of transient receptor potential vanilloid type-1 (TRPV1), a nonselective cation channel activated during hypertonicity-induced neuronal shrinking^7-10^. Interestingly, the microtubules (MTs) components of the cytoskeleton play a vital role in this mechano-transduction process, forming a complex scaffold with TRPV1 that is activated during cell shrinking and bidirectionally influences osmosensory gain^11-13^.

Recent electrophysiological studies have evidenced the generation of self-sustained oscillations in brain MTs^14,15^. Empirical mode decomposition (EMD) analysis of these oscillations^16^ indicates their underlying contribution to identifiable brain waves observed by EEG and LFP^17-^. These studies underscore the capacity of brain MTs to act as intracellular oscillators observed in various mammalian species. The study by Gutierrez et al.^26^ further supports the idea that MT-driven oscillations may serve as the intrinsic oscillator of the brain. A growing body of evidence suggests a crucial role for MTs in the dynamic regulation of neuronal activity and underscores their significance in understanding the fundamental mechanisms underlying brain function and dysfunction. Thus, intrinsically generated MT oscillations may regulate task specific neuronal function with implications for the osmoregulation of neurons possessing unique MT architectures as the hypothalamic MCNs.

To explore this phenomenon, we investigated the impact of dehydration on the electrical activity of the MT-based cytoskeleton of the brain in adult rats. We observed electrical oscillations in SON samples that changed their properties in water deprived animals. Our findings revealed that dehydration alters MT-based electrical activity in the SON, a key region involved in fluid balance regulation, a phenomenon that was not observed in samples of the cerebral cortex. Thus, brain MTs respond to water deprivation by modulating electrical oscillations, thereby contributing to the organism’s homeostatic response through a novel cytoskeletal signaling mechanism.

## Results

### Electrical activity of adult rat SON tissue

To obtain electrical information from the SON, tissue matter was obtained as described in the Materials and Methods from control and DH animals^27^. Samples were processed, and kept frozen at -20oC until the experiment, when 1-2 mm size samples were defrosted into normal (high NaCl) saline solution. Electrical recordings were conducted as in Gutierrez et al.^26^, and the companion manuscript^27^ (Fig. 1). Experiments were conducted under symmetrical (pipette and bathing solution) conditions, in a Ca^2+^-free “intracellular-like” solution containing high KCl (140 mM) and 1 mM EGTA. Current-to-voltage relationships were obtained for control and DH SON (Fig. 1B), where sigmoidal curves from DH SON area were observed with much higher values for the control samples.

**Fig. 1:**
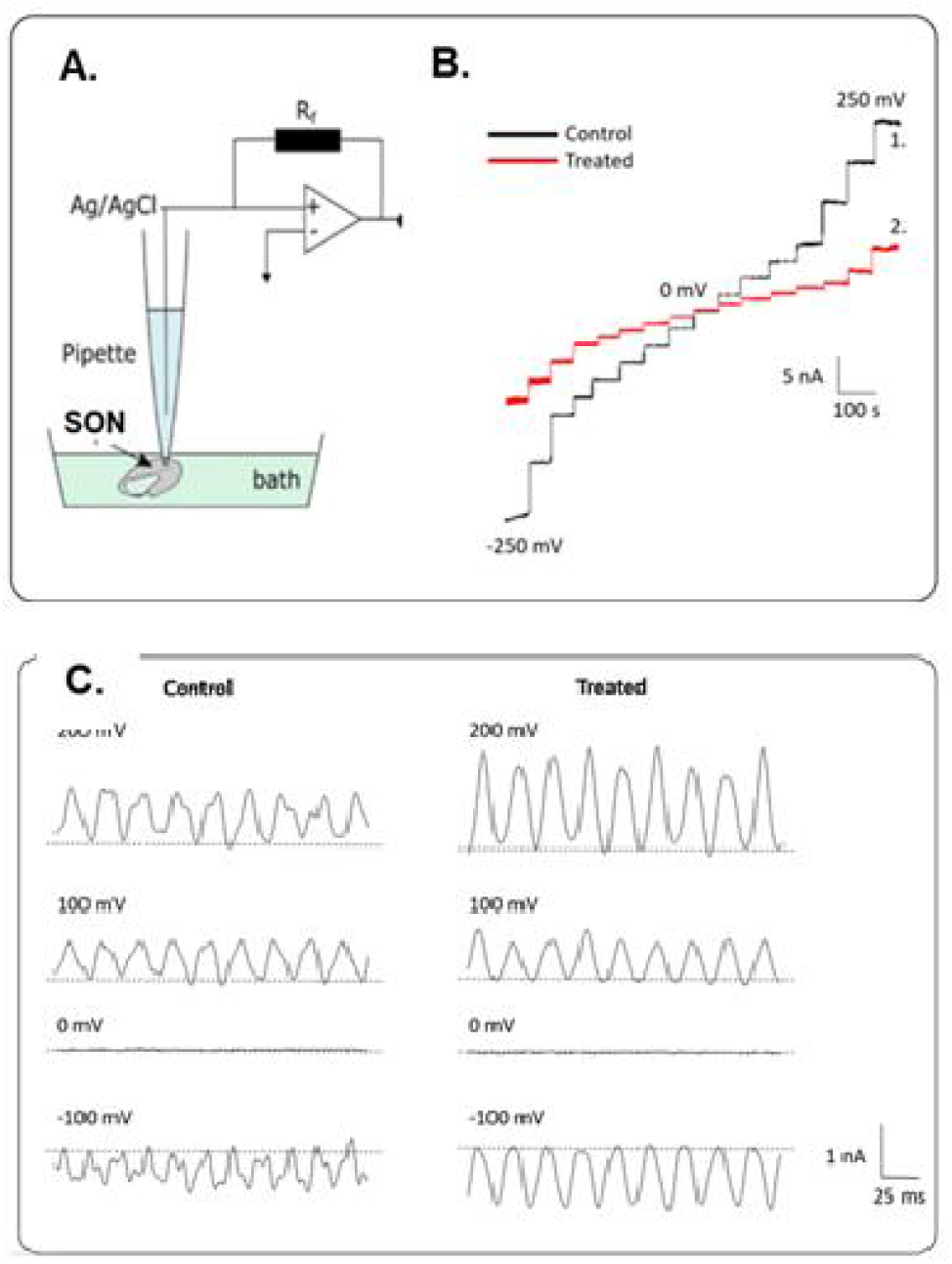
Experimental setup and data analysis. **A**. Technical outline showing the loose-patch data acquisition system. **B**. Current obtained at several voltages from -250 mV to 250 mV from control (Black line) and dehydrated (red line) SON. **C**. Representative tracings of electrical currents obtained at different voltages (as indicated in the panels) for SON samples from either control or DH animals. Please note that while the overall conductance was much larger in the control sample (B), the oscillatory response has a higher amplitude in the samples from DH animals.

Electrical activity of rat brain tissue in the presence of KCl in the bath solution. Rat brain tissue samples (n = 43) were tested with a patch clamp amplifier (Fig. 1A-B), as recently reported^27^, and a pipette electrode filled with an intracellular type of saline, as previously reported^26^. The voltage-clamp patch pipette was used to assess local field potentials (LFP) in the form of electric currents at the location of the pipette under symmetrical ionic conditions. The tip resistance of the patch pipette was 5.76 ± 1.38 MΩ (n = 32) under symmetrical saline (identical KCl concentration in pipette and bath) conditions. The tip conductance was first measured in saline solution, which was highly linear (data not shown) and devoid of any oscillatory behavior, except for the line frequency at ∼50 Hz (Fig 1C), while distinct patterns of frequencies are observed in the SON-patched power spectra (Fig. 1D).

Voltage-clamped SON samples displayed spontaneous, self-sustained electrical oscillations that responded to the magnitude and polarity of the electrical stimulus (Fig. 2A) in both conditions. In samples obtained from DH-treated rats, the currents were more prominent at high voltages (Fig. 2A,B,D). Current-to-voltage relationships were obtained for the average of electrical oscillations (Fig. 2C) from SON samples under control and DH-treated conditions to explore the electrical features of the samples further. The current-voltage relationships obtained without any correction were almost linear in both conditions and showed more resistance for treated samples (slopes, Fig. 2C). However, a closer inspection of the tracings showed that for the DH-treated samples, the oscillations were more prominent at high voltages (>200 mV). The magnitude of the difference between the maximal and the minimal currents under each condition disclosed a highly nonlinear response in the DH-treated samples not observed under control condition (Fig. 2C). The nonlinear conductance was clearly more prominent for the DH-treated samples than the controls, as observed for pooled data in the range of 200-250 mV (Fig. 2D). Three-dimensional phase-space portraits showed limit cycles, evidencing mono-periodic oscillatory behavior. An example at 250 mV is shown in Fig. 2E.

**Fig. 2:**
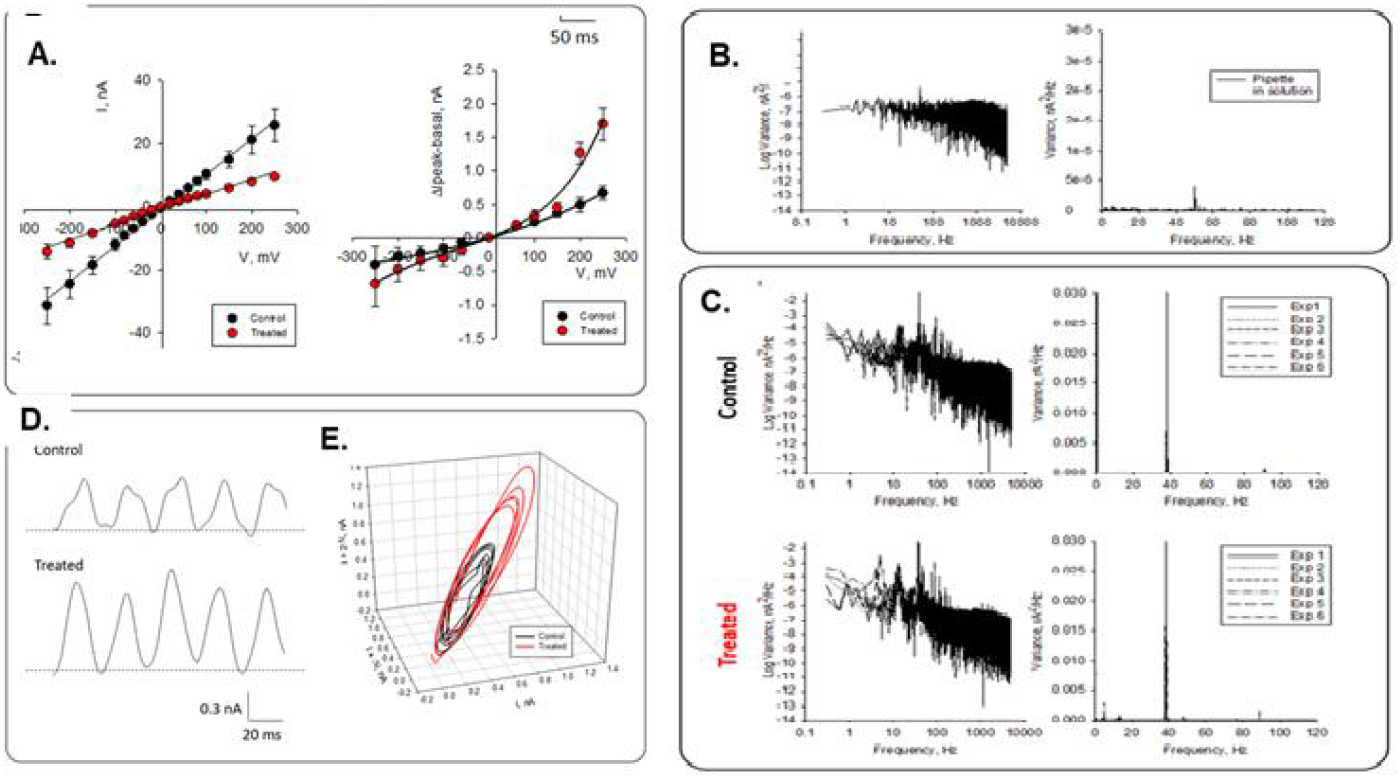
Electrical oscillation from SON samples. ***A***. Representative current-to-voltage relationships were obtained for the difference between the peak and the basal current for either control (black circles, *n* = 8) and treated (DH, red circles, *n* = 6) conditions. Solid lines represent the average conductance of each preparation. **B**. The frequency spectrum of electrical unfiltered current tracing through the pipette in saline solution shows the frequency peak of the power line expected at 50 Hz. **C**. Top. The spectra of SON in control conditions showed a prominent peak at 39 Hz and a minor peak at 90 Hz (*n* = 6). Bottom. Spectra of SON in treated conditions showed a prominent peak at 39 Hz and minor peaks at around 5 and 10 Hz (*n* = 6). FFT values were conducted from tracings obtained at 100 mV. **D**. Expanding tracings indicated (1 and 2) in Fig. 1B comparing the currents at 250 mV in control and treated conditions. **E**. 3D phase portrait for control (black line) and DH-teatred (red line) conditions, in the presence of KCl in the pipette and bathing solutions.

To further explore the frequency of the obtained oscillations, we analyzed the traces using Fourier transformation. Fourier spectra showed a fundamental frequency of ∼38 Hz and minor peaks of ∼5, ∼17, ∼79, and ∼93 Hz after impaling control and DH-treated tissues. Interestingly, the graphs show that the variances of the minor peaks were different for both the tissue and the DH-treatment (Fig. 3).

**Fig. 3:**
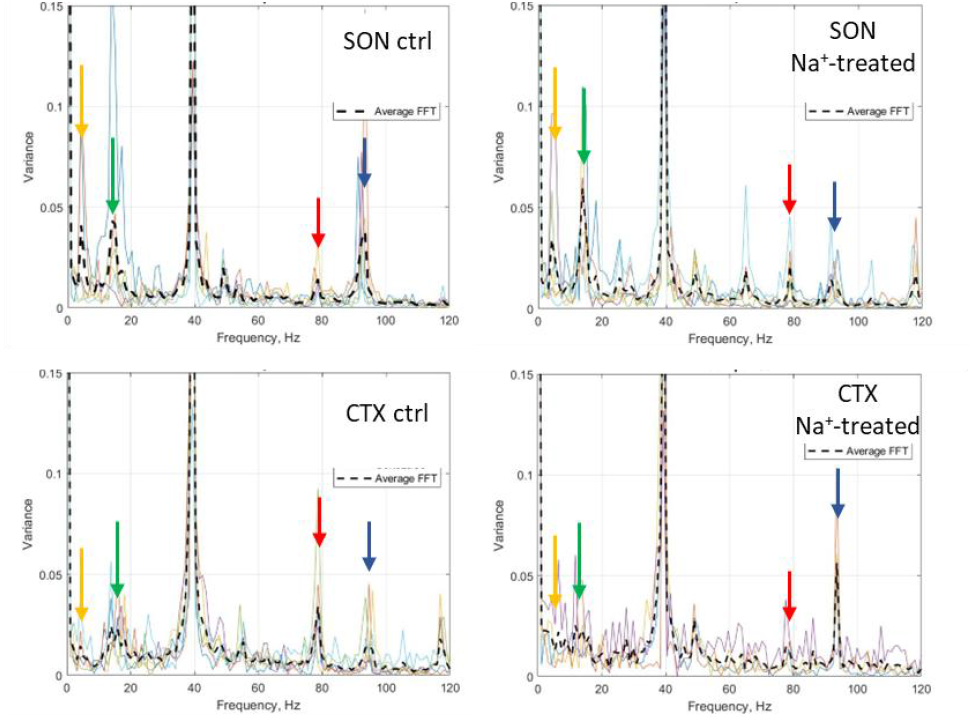
FFT distribution of oscillatory frequencies from control and DH-treated CTX and SON. **Top**. Fourier spectra for SON tracings in control (ctrl, left, *n* = 6)) and DH (right, *n* = 6) conditions. **Bottom**. Fourier spectra for CTX tracings in control (left, *n* = 6) and DH-treated (right, *n* = 6) conditions. Distribution values from individual experiments are shown in different colors and averages in black dashes. Colored arrows indicate frequencies of interest.

Empirical Mode Decomposition of rat brain samples. To explore the intrinsic differences between the rat CTX and SON-generated electrical oscillations, in control and DH conditions, we subjected representative electrical signals to EMD analysis, as recently reported^16^. CTX samples showed 7 to 9 IMFs (control n = 6, DH-treated n = 4), while the SON displayed 8 IMFs in control conditions and 6 to 8 IMFs for DH-treated tissue (control n = 6, Na+-treated n = 6). Fig. 4 depicts an example of control and DH SON (CTX EMD in^27^.

**Fig. 4:**
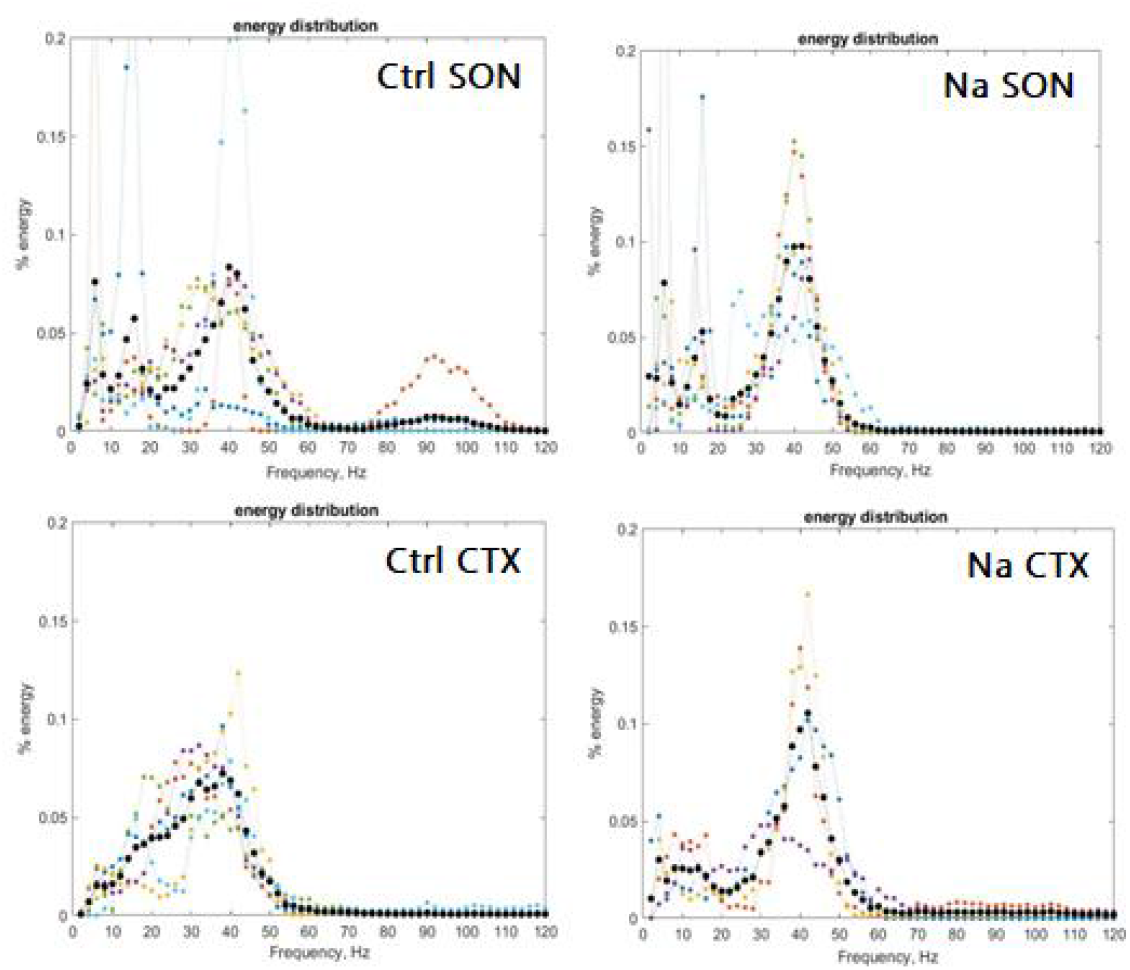
Energy distribution of oscillatory frequencies from control and DH-treated CTX and SON. Distribution values from individual experiments are shown in different colors and averages in black circles.

Further, these strategies confirmed other frequencies beyond the fundamental one, and differences between samples in five fundamental frequency ranges: 3-8, 10-20, 33-44, 70-83 and 83-100 Hz. The energy per frequency range was calculated as a percentage of the reduced area under the curve (%RAUC, Table 1, see Supplementary Material) based on the Fourier transforms to obtain information about the frequency density. The energy contributed by each IMF was further explored using the HHT and CWT decompositions (Fig. S1, Supplementary Material). The CWT decomposition method displayed higher frequencies not observed in the HHT plots. However, no details could be gathered from each contribution by these methods.

Effect of dehydration adaptation on the electrical oscillations of the cortex and SON. To explore the possible selective electrical oscillatory response of the SON in DH animals, we also chose another region of the brain, which we have electrically characterized, the brain cortex^27^.

Energy distribution of peak frequencies between CTX and SON. We explored the oscillatory currents in more detail to identify specific energy contributions from the various frequency ranges: 3-8 Hz, 10-20 Hz, 33-44 Hz, 70-83 Hz, and 83-100 Hz. Under control conditions, we observed energy differences of the CTX and SON in the range of 3-8 Hz, where SON samples were 277% higher than CTX (7.25 vs. 1.92, p = 0.016) (Fig. 5A). The fundamental frequency range of 33-44 Hz was highly similar among groups. However, in the ranges 70-83 Hz and 83-100 Hz, the SON samples were 38.8% (2.83 vs. 7.29, p = 0.005) and 40.5% (10.62 vs. 4.30, p = 0.029) lower than CTX, respectively (Fig. 5). This difference agrees with the observed frequency differences between cortex and hippocampus^27^.

**Fig. 5:**
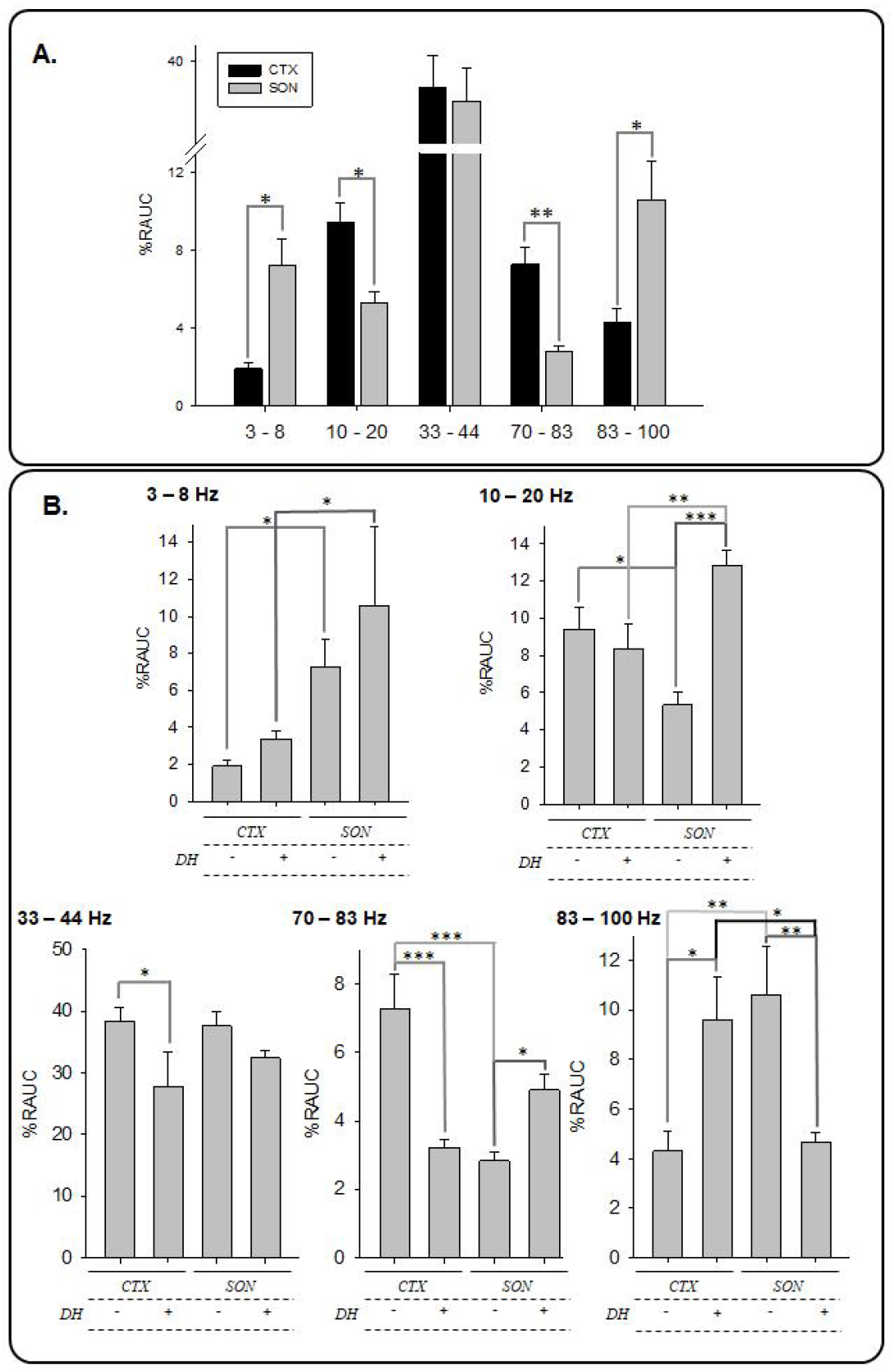
Energy distribution of frequency ranges from control and DH-treated CTX and SON. **A**. Comparison of distribution values for the identified frequency ranges of the two brain regions, cortex (CTX), and supraoptic nucleus (SON). **B**. Differences of frequency ranges for the control (Ctrl) and DH-treated (DH) CTX and SON samples. Statistical differences are marked as * for *p* < 0.05, ** for *p* < 0.01, and *** for *p* < 0.001 for the various frequency ranges.

DH treatment showed a tissue-specific response at identifiable frequency peaks, including the 10-20 Hz range, where SON responded with a 141% increase in signal (12.87 vs. 5.33, p < 0.001), while we found no response in CTX (9.41 vs. 8.38, p = NS). The DH response did not impact the fundamental frequency range (33-44 Hz, p = NS), however in the 70-83 Hz, while CTX responded with a substantial signal reduction (7.29 vs. 3.21, p < 0.001), SON doubled (2.83 vs. 4.89, p = 0.032). Conversely, the 83-100 Hz range showed a substantial increase in the CTX (9.60 vs. 4.30, p = 0.029) with a reduction in the SON peak (10.62 vs. 4.64, p <0.005).

Electron microscopy of brain tissue samples. To explore the morphological correlates to the electrical properties of the brain tissue, histological samples were obtained from either SON or CTX from the same samples recorded and analyzed by electron microscopy (Fig. 6). Samples were negatively stained, and images were obtained at 10 kV. Longitudinal tracks of MTs that showed different inter-MT distances between control and DH SON samples were observed, with Ctrl 40.59 ± 0.67 nm (n = 11), vs. DH samples of 32.58 ± 2.84 (n = 11), p 0.0125. Cortex samples did not show any statistical differences between control and dehydrated conditions (data not shown). Evidence of functional differences in the brain MTs of different locations is further observed in the accompanying manuscript, where special differences in intra-axonal MT tracks were not observed between different brain regions^27^.

**Fig. 6:**
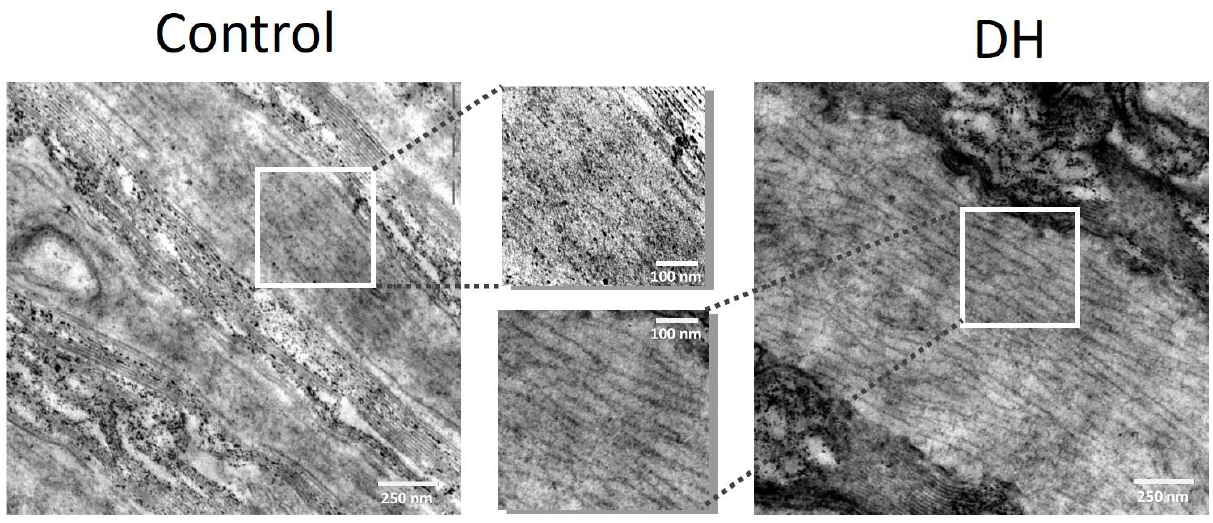
Electron microscopy of SON samples. Left, control condition, and Right, dehydrated (DH)- condition. Insets (white squares) are shown on the Center, evidencing the density of MT distributions.

The inter-MT distance of 40.6 ± 0.67, n = 11 under control conditions, was reduced to 32.6 ± 2.80 nm, n = 11 (p = 0.0125), after dehydration.

## Discussion

The MT cytoskeleton is a dynamic network of cylindrical structures, which gives cells their shape, organizes the cell’s internal structure, and plays critical roles in various cellular processes. MTs provide structural support to maintain and enable changes in cell shape, and are implicated in the transport of organelles, vesicles, and proteins within the cell. MTs are crucial in mitosis and meiosis, where they form the mitotic spindle. MTs are also implicated in cell sensory and motility functions, where the axoneme of cilia and flagella may act as electrical antennas^28^. Thus, the special arrangements of MT assemblies may be essential for distinct intracellular electrical signaling. Evidence of this is also suggested by the uniquely interweaved scaffold of intrasoma MTs in osmosensory neurons of the SON, where they physically interact with the carboxy terminus of TRPV1 at the cell surface to provide a pushing force that drives channels activation during osmotic shrinking^13^. MTs are thus an essential component of the vital neuronal mechanotransduction apparatus that allows the brain to monitor and correct body hydration^13^.

The present study investigated the hypothesis that a specific MT organization in SON osmosensitive neurons responds, through its electrical oscillation, to an in vivo osmotic challenge. Acute dehydration from water deprivation produced an osmotic stimulation that gates several homeostatic responses.

A companion study explored the intrinsic electrical activity of brain MTs in identified brain regions, including the cortex and hippocampus^27^. In-depth analyses of brain tissue from the hippocampus and neocortex identified specific electrical properties of MTs. MTs act as biomolecular transistors, amplifying electrical signals across their intramolecular junctions^29^ (whose ionic conductivity due to nanopores formed by their lateral arrangement of protofilaments is sensitive to the antineoplastic paclitaxel^30^ that inhibits these oscillations in a voltage-dependent manner. The present studies revealed spontaneous electrical oscillations with distinct characteristics and differences in mean conductances between brain regions.

The evidence shown in this manuscript demonstrates the presence of spontaneous oscillations in the SON, with significant differences in amplitude and intrinsic frequencies between control and the DH-treated animal samples. These findings agree with the presence of an intrinsic oscillator possibly contributed by the electrical activity of neuronal MTs. This was confirmed by comparison of the DH-response in other non-osmotically sensitive brain regions such as the neocortex which did not significantly change its response to in vivo water deprivation induced dehydration. In osmosensitive neurons, MTs extend to the cell surface and interact closely with the plasma membrane, binding to TRPV1 channels^13^. A disruption of the TRPV1-MT interaction blocks the excitation of ONs induced by mechanical or osmotically induced shrinking, emphasizing the importance of this structure-function interaction^13^. The dense MT network in ONs equilibrates compression forces along the cell surface during volume changes induced by osmotic perturbations, potentially optimizing the monitoring of changes in MT scaffold compression and preserving the ONs structural integrity.

In agreement with our findings, studies carried out with synchrotron small-angle organization (SAXS) and transmission electron microscopy (TEM), revealed that changes in osmotic pressure and/or in the valence of cations influence the organization/ interaction and spacing of MTs bundles^31,32^. It is possible that this osmotic effect on the MT, which alter the nature and magnitude of interactions, also influences the electrical oscillations of MTs in the face of changes in osmolarity induced by dehydration. In summary, our present results revealed that an acute dehydration, induced by 24 h water deprivation, alters MT-based electrical activity in the SON, a critical area involved in fluid balance regulation. However, this phenomenon was not observed in samples of the cerebral cortex. Thus, brain MTs respond to dehydration by modulating electrical oscillations in osmosensitive/osmo responsive neurons, thereby contributing to the organism’s homeostasis through a novel cytoskeletal signaling mechanism.

Interestingly, our findings are in general agreement with previous reports on the effects associated with dehydration and hangovers, which significantly affect brain wave activity, as reflected in EEG recordings, despite the fact that their patterns differ based on the cause of fluid loss. Dehydration from exercise or lack of fluid intake is associated with increased slow-wave activity, including theta and delta waves, along with a reduction in alpha power, which correlates with cognitive decline and mental fatigue^33,34^. Further, dehydration leads to a reversible decrease in beta wave activity, impairing attention and cognitive performance^35,36^. Increased cortical excitability is another hallmark of dehydration, which can be manifested as heightened EEG activity and impaired cognitive regulation^37^. In contrast, alcohol-induced dehydration during hangovers shows a distinct EEG pattern, characterized by elevated alpha and theta activity, which is reflected in impaired cognitive function and sensory processing^38,39^. Rehydration normalizes these EEG patterns, improving mental clarity and restoring cognitive performance^40,41^.

We have recently described how the trancriptome, proteome and the phosphoproteome of the SON are affected by dehydration^42,44^. Interestingly, pathway analysis revealed that post-translational protein phosphorylation modifications in the SON mediate cytoskeleton remodeling. These events involved the cytoskeletal proteins DBNL, HSP90AB1, MAP1A, MAP1B, MAPT, ribosomal protein S3 (RPS3), Stathmin 2 (STMN2), and also proteins involved in cytoskeletal protein binding, namely ANK2, MAP2, STMN1. All these proteins were hyperphosphorylated, with the exception of STMN2 that was hypophosphorylated and MAP1B and MAPT that were both hyper and hypophosphorylated at different residues. Whether these dynamic phosphorylation events are involved in the electrical phenomenon described here remains to be determined.

## Conclusions

The present study demonstrated that the electrical oscillations generated by MTs in the SON of the adult rat are modified after dehydration. Thus, the physiological response of this brain region includes a distinct electrical response of the cytoskeleton, which is likely associated with morphological parameters such as inter-MT distance that affect the pattern of electrical oscillations.

## Methods

### Animals

We used 12 adult male Wistar rats, born and reared in the breeding colony at Instituto Ferreyra (INIMEC-CONICET-UNC, Córdoba, Argentina). Please note that the number of experiments under any given condition (as indicated in the Results section) may not reflect the total number of animal in their respective group. Animals weighing 250–300 g were housed singly in metabolic cages with free access to a normal diet (Purina Rat Chow) and distilled water for three days of adaptation. Room lights were on for 12 h/day, kept at ∼23ºC. All experimental protocols were approved by our institute’s appropriate animal care and use committee under protocol # 016/2021, following the guidelines of the International Public Health Service Guide for the Care and Use of Laboratory Animals (NIH Publications No. 8023, revised 1978). We complied with the ARRIVE guidelines.

### Water derivation protocol

after the adaptation period, animals were randomly divided in two groups: one remained in the same conditions as in the habituation period (euhydrated: control group) and another was maintained for the next 24 h with free access to standard chow but no access to fluids (DH, dehydrated group). At the end of 24 h, rats were euthanized by decapitation without previous anesthesia since the general anesthetics may change MT electrical activity (Cantero, unpublished data). Brain dissection were performed as follows.

### Brain areas dissection

Immediately after decapitation, the brains were collected and placed in an ice bed for dissection. Serial coronal sections of 300 μm from CTX (bregma -0.80 to - 1.80mm - cingular cortex Cg1 and Cg2 and part of primary and secondary motor M1 and M2, respectively) and SON (bregma: -2.12 to -4.16 mm, Paxinos, and Watson, 2007) were obtained from the brain using a vibratome. Identified brain regions were taken using stainless-steel needle punches of 2 mm. The extracted area included mainly the. Brain samples were washed and kept in an extracellular solution containing, in mM: 135 NaCl, 0.5 KCl, 1 MgSO_4_, 1.5 CaCl_2_, 1 EGTA, 10 HEPES, and pH adjusted to 7.23). The sections were frozen in this solution at -20oC until the time of the experiment. Note that fresh samples were also tested that had only been refrigerated until the time of the experiment, with similar results. Tissue freezing did not affect the electrical activity observed in the present study and was thus used throughout the experiments described therein.

### Electrophysiology

Electrical recordings from brain tissue were obtained with the loose-patch-clamp configuration, as recently reported for Apis mellifera brain^26^. Briefly, command voltages (V_cmd_)were applied inside the brain matter of the exposed tissue. Electrode setup was similar to patch clamping with pipettes filled with a solution containing, in mM: KCl 140, NaCl 5, EGTA 1.0, and HEPES 10, adjusted to pH 7.18 with KOH. Approximately 2 mm size tissue samples were added to the dry surface of the patch clamp chamber, letting it rest for 5 min before adding 400 μl of saline solution. Experiments were conducted under symmetrical conditions, with an “external” bathing KCl solution containing the same as the pipette^14^. Electrical recordings were conducted with a miniaturized patch-clamp amplifier, ePatch, from Elements (Cesena, Italy) with a recording range between ±200 nA, voltage stimulus range ±500 mV, and a maximum signal bandwidth of 100 kHz. Patch pipettes were made from soda lime 1.25 mm internal diameter capillaries (Biocap, Buenos Aires, Argentina) with tip diameter of ∼4 μm and tip resistance in the order of 5-15 MΩ. Voltage clamp protocols only included step-wise holding potentials (gap-free protocol) from zero mV. Electrical signals were acquired at 10 kHz and stored in a computer with the EZ patch software 1.2.11 (Elements, Cesena, Italy). Sigmaplot Version 11.0 (Jandel Scientific, Corte Madera, CA) was used for statistical analysis and graphics. Power spectra of unfiltered data were obtained by the Fourier transform subroutine of Clampfit 10.0.

### Transmission electron microscopy

Samples of cortex or the hippocampus were processed following the method described previously^45^ with slight modifications for brain tissue. Briefly, tissue small sections (10-20 mm^3^) were fixed with Karnovsky’s fixative (a mixture of 2.66% w/v paraformaldehyde and 1.66% w/v glutaraldehyde in 0.1 M phosphate buffer, pH 7.2), overnight at 4°C. After fixation, cells were washed with 0.1 M phosphate buffer and embedded in 1.2% agar to form easy-to-handle sample blocks. The samples were post-fixed in 1% with osmium tetroxide for 2 hours at 4°C. After washing with phosphate buffer, they were block stained with 2% uranyl acetate for 30 minutes at room temperature in the dark. Subsequently, the samples were dehydrated in a graded ethanol series (50%, 70%, 90%, and 100%), followed by 100% acetone. Samples were then infiltrated and embedded in an acetone-SPURR resin sequence (SPURR resin; Ted Pella, Inc.). They were transferred to embedding plates and supplemented with fresh embedding medium. Polymerization was carried out at 60°C for 24 hours. After polymerization, thin sections were cut with and Powertome XL Ultramicrotome (Boeckeler Instruments, Inc.) and mounted on uncoated 200-mesh copper grids (Ted Pella). All grids were examined with a transmission electron microscope (Zeiss LIBRA 120; Carl Zeiss AG, Germany) at 80 kV, at the Electron Microscopy Core Facility and Research Center (Centro Integral de Microscopía Electrónica-CIME-CONICET-UNT).

### Drugs and chemicals

All reagents were obtained from Sigma-Aldrich (St. Louis, MO USA) unless otherwise stated. Wherever indicated, the microtubule stabilizer paclitaxel (Taxol Equivalent, Invitrogen™, P3456) was prepared as per the manufacturer’s recommendations (10 mM in DMSO) and added at the indicated concentrations.

### Empirical Mode Decomposition (EMD) Analysis

As previously reported, the EMD method was performed^16^ to analyze the data in the Time-Frequency (TF) domain. The original signal was decomposed into Intrinsic Mode Functions (IMFs), waveform functions with only one frequency. The signals were filtered by a Gaussian filter at 200 Hz, Notch for electrical noise (50 Hz), and its harmonics using Clampfit 10.7 (Molecular Devices). Preconditioned data was parametrized using a custom function of Matlab 2019a (Mathworks, Natick, Massachusetts), and then the Matlab “emd” function was performed to obtain the signals IMFs. Each IMF coefficient (and thus its frequency) was obtained with the Matlab “Curve Fitting Tool.” The Hilbert-Huang Transform (HHT) graphics show the EMD, followed by the Huang transformation^46^ to present the IMFs with an energy-frequency-time distribution. Graphics were obtained using the Matlab function “htt”.

### Energy Calculations

To obtain the Continuous Wavelet Transform (CWT) that decomposes the original time signal (“mother wavelet”), and the energy implied on each daughter wavelet, we used, as recently reported^16^ the “cwt” Matlab function, which uses the analytic Morse wavelet with standard parameters. The CWT frequency-time plot helper was used for the graphics. Relative energy was estimated using the Fourier spectrum, calculating the area under the curve (AUC) within the range, as reported for electroencephalographic (EEG) records^47,48^. The original Fourier spectrum had a maximum frequency of 5000 Hz in all cases. A reduced frequency range was then selected to provide a Reduced Area Under the Curve (RAUC), from 0 to 140 Hz, that resembles the use of Spectral Edge Frequency (SEF) in the analysis of EEG^49^. The frequency ranges were the same as reported^16^ for comparison. The percentage of energy involved in each IMF per frequency peak was also calculated and graphed using each IMF instant frequency and energy with a 2D histogram Matlab custom function.

### Statistical analyses

Mean IMF frequency was expressed as mean and standard deviation (SD). The % RAUC was represented with mean value and standard error (SE). % RAUC per range was compared between samples using t-test or Mann-Whitney if variable normality (Shapiro Wilk) or homoscedasticity (Levenne test) failed. For statistics and graphics, SigmaPlot 11.0 software was used. Inter MT distances from TEM images were analyzed as a quantitative variable, observing mean plus minus standard error (t test), for n measurements of two different photographs. Statistical significance was set at p < 0.05.

## Acknowledgements

MdRC and HC wish to acknowledge partial funding of the present study to the Ministerio de Ciencia, Técnica, e Innovación, Argentina (PICT 0050, 2021), and CONICET, PIBAA (0495). DM thanks the Medical Research Council for generous support (MR/W028999/1).

## Author contributions statement

The authors contributions were as follows, AG, MdRC, and HFC conceived the experiments. CP and AG conducted all animal experimentation and preparation of the samples MdRC conducted the electrophysiology experiments, curated the data, and analyzed the results. DM and ASM further analyzed the results, and VHA conducted the electron microscopy studies, and HFC, MdRC, AG, ASM, and DM, designed the experiments, and prepared the draft. All authors read the manuscript.

## Additional information

Competing interests. The authors declare no competing interests

## Supplemental Material

**SM-Fig.1:**
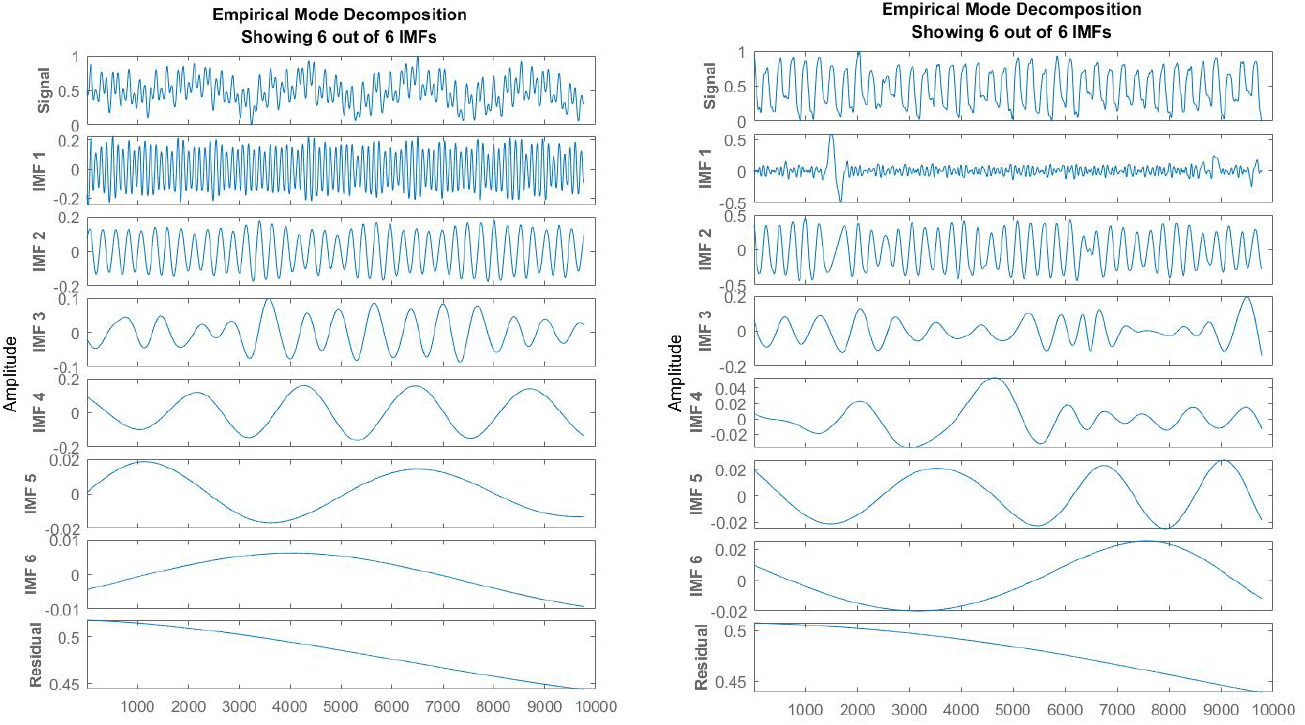
Empirical Mode Decomposition (EMD) of SON tracings. The EMD displayed 6 to 8 IMFs in control (left, n = 6) and DH (right, n = 6) conditions.

## Notes

### Competing Interest Statement

The authors have declared no competing interest.

### Summary of Updates

We corrected one reference of the list

